# Bacteriophage inhibits murine norovirus replication after co-incubation in RAW 264.7 cells

**DOI:** 10.1101/2021.03.09.434703

**Authors:** Lili Zhang, Khashayar Shahin, Ran Wang

## Abstract

Bacteriophages (phages) are viruses of bacteria. Despite the growing progress in research on phage interactions with eukaryotic cells, our understanding of the roles of phages and their potential implications remains incomplete. The objective of this study was to investigate the effects of the *Staphylococcus aureus* phage vB_SauM_JS25 on murine norovirus (MNV) replication. Experiments were performed using the RAW 264.7 cell line. After phage treatment, MNV multiplication was significantly inhibited, as indicated by real-time quantitative PCR (RT-qPCR) analysis, western blot, and immunofluorescence assay. Furthermore, we demonstrated the transcriptional changes in phage–MNV co-incubated RAW 264.7 cells through RNA sequencing (RNA-Seq) and bioinformatic analysis. Our subsequent analyses revealed that the innate immune response may play an important role in the restriction of MNV replication, such as the cellular response to IFN-γ and response to IFN-γ. In addition, the gene expression of *IL-10, Arg-1, Ccl22, GBP2, GBP3, GBP5*, and *GBP7* was proven to increase significantly by RT-qPCR, showing a strong correlation between RT-qPCR and RNA-Seq results. Furthermore, phage treatment activated guanylate binding proteins (GBPs), as revealed by RT-qPCR analysis, western blotting, and confocal microscopy. Taken together, these data suggest that the phage affects the innate response (such as the IFN-inducible GTPases and GBPs), reflecting their direct antimicrobial effect on the membrane structure of the MNV replication complexes, and therefore, exerts an antiviral effect *in vitro*. Collectively, our findings provide insights into the interactions of immune cells and phages, which improve our understanding of the actual role and potential of phages.

## Introduction

Bacteriophages (phages) are natural predators of bacteria. So far, the main application of phages has been the treatment of bacterial infections. In fact, phages can interact with eukaryotic cells (especially cells of the immune system). However, our understanding of the roles of phages and their potential implications remains incomplete. To date, there have been conflicting reports on the potential antiinflammatory action of administered phages (Van Belleghem, et al., 2017, Zhang, et al., 2018). Most *in vivo* studies to date describing lytic phage-induced antiinflammatory effects report the simultaneous clearance of bacterial infections, which may in fact be the driving force in relieving inflammation (Zimecki, et al., 2009, Miedzybrodzki, et al., 2017).

Apart from antibacterial activity, phages can also exert antiviral activity (Miedzybrodzki, et al., 2005, Sweere, et al., 2019). For instance, the internal phage protein phagicin showed activity against vaccinia virus, by interfering with the synthesis of viral DNA (Meek and Takahashi, 1968). Another line of research took into consideration the antiviral action of phages via nucleic acids, both purified dsRNA and ssDNA, revealing activity against herpes simplex virus, duck hepatitis B virus, and vaccinia virus (Feldmane, et al., 1977, Vales, et al., 1991, Iizuka, et al., 1994, Mori, et al., 1996). The third proposed model of interaction is based on the competition for beta3 integrins between the Lys-Gly-Asp (KGD) motif of *Escherichia coli* phage T4 and viruses utilizing integrins as a portal of entry (Dabrowska, et al., 2004). In addition, both *E. coli* phage T4 and *staphylococcal* phage A5/80 significantly reduced the expression of the human adenovirus type 5 *(HAdV-5*) gene, while A5/80 does not possess the KGD motif. The proposed mode of antiviral action is related to a DNA-binding protein (Przybylski, et al., 2019). In addition, there are a variety of intracellular innate immune mechanisms, which may be initiated in phage–virus interactions during viral infection (Van Belleghem, et al., 2018).

In this work, we systematically evaluated the effects of a *Staphylococcus aureus* phage on the course of infection by a pathogenic virus. Murine norovirus (MNV), which is closely related to human norovirus, is an RNA virus that can prove lethal in mice with impaired innate immunity. The main objective of the present study was to investigate the effects of phage vB_SauM_JS25 on the replication of MNV after direct induction of immune cells. This study expands our understanding of the role of phages.

## Materials and Methods

### Bacterial strain, bacteriophage stock, cell line, murine norovirus, and growth conditions

*S. aureus* strain ATCC 6538 was grown in tryptic soy broth (Qingdao Hope Bio-technology Co., Ltd., Qingdao, China) at 37°C with shaking overnight and used to propagate and quantify phage vB_SauM_JS25. Phage lysates were further purified using cesium chloride (CsCl) centrifugation to remove toxins, as described previously (Zhang, et al., 2018). The phages were ultrafiltered through a cellulose membrane (Millipore, Billerica, MA) with a nominal molecular mass limit of 100 kDa to remove excess CsCl. A blank sample was also used from bacterial lysates. The bacterial cells were disrupted under ultrasonication and centrifuged at 8,000 × g for 20 min. The phage supernatant was filter-sterilized using a 0.22 μm filter to yield a bacterial cell-free lysate. Compared to phage ultracentrifugation in CsCl gradients, the same phase was collected using a syringe, following the protocol of phage purification. Cells of the murine macrophage cell line RAW 264.7 were maintained in Dulbecco’s Modified Eagle’s Medium (DMEM) supplemented with 10% fetal calf serum (Sigma-Aldrich, MO, USA) and grown at 37°C with 5% CO_2_. The MNV strain Nanjing 1608 (GenBank accession no.: KX987840), was propagated in RAW 264.7 cells as previously described (Ma, et al., 2020).

### Measurement of endotoxin and *Staphylococcal* enterotoxins

The phage endotoxin concentration was measured using the chromogenic Limulus Amebocyte Lysate ToxinSensor^™^ kit (Genscript Corporation, NJ, USA, < 0.01 EU/ml) and the *Staphylococcal* enterotoxin (SET, SEA to SEE) concentration was measured using the Evergreen TREA^™^ SET total assay kit (Evergreen Sciences Inc., CA, USA, < 0.25 ng/ml) according to the manufacturer’s protocols.

### Effect of bacteriophage vB_SauM_JS25 on MNV multiplication

To assess the capacity of phage vB_SauM_JS25 to affect MNV multiplication in RAW 264.7 cells, which are susceptible to MNV infection, the cells were seeded and incubated overnight at 37°C with 5% CO_2_. When the cells reached 80–90% confluence, they were washed and inoculated with MNV (at a multiplicity of infection [MOI] of 1) at 37°C. After 1 h, the cells were washed with PBS to aspirate the inoculums and purified phage vB_SauM_JS25 was added to the cells at various titers (10^3^ to 10^8^ PFU/mL). At the indicated time points post-phage treatment, the cells were collected and the effect of phage vB_SauM_JS25 on MNV replication was detected by real-time quantitative PCR (RT-qPCR).

### Cell viability assay

RAW264.7 cells were seeded into 96-well plates and treated with phage, MNV, or phage plus MNV. At the indicated time points, cell viability was assessed using a Cell Counting Kit-8 (CCK-8, Dojindo, Japan) following the manufacturer’s protocol. The cytotoxic concentration was calculated from three independent experiments and is expressed as the mean ± SD.

### Viral binding test on MNV of bacteriophage vB_SauM_JS25

The viral binding test on MNV was performed as previously described (Li, et al., 2016). RAW 264.7 cells were seeded into 24-well plates at a density of 5 × 10^5^ viable cells per well. On the following day, MNV (MOI, 1) and phage vB_SauM_JS25 (10^8^ PFU/mL) were added and the plates were incubated for 1 h at room temperature. Afterward the inoculum was aspirated and the cells were washed by PBS for three times. The binding effect of phage vB_SauM_JS25 on MNV replication was analyzed by RT-qPCR.

### Real-time qPCR

Total RNA was extracted from cellular samples using an RNA kit (Omega Biotek, GA, USA) in accordance with the manufacturer’s instructions. Then, 500 ng of total RNA was used in 10 μL PrimeScript^™^ RT Master Mix (TaKaRa, Tokyo, Japan) as described in the manufacturer’s instructions. Next, the cDNA was used as a template for RT-qPCR, which was performed on a LightCycler 480 (Roche Diagnostic, IN, USA). The PCR primers have been described previously(Biering, et al., 2017, Ma, et al., 2020) and are presented in Table 1. Relative gene expression was determined using the 2^-ΔΔCt^ method by normalization with the *GAPDH* reference gene.

**Table 1.**
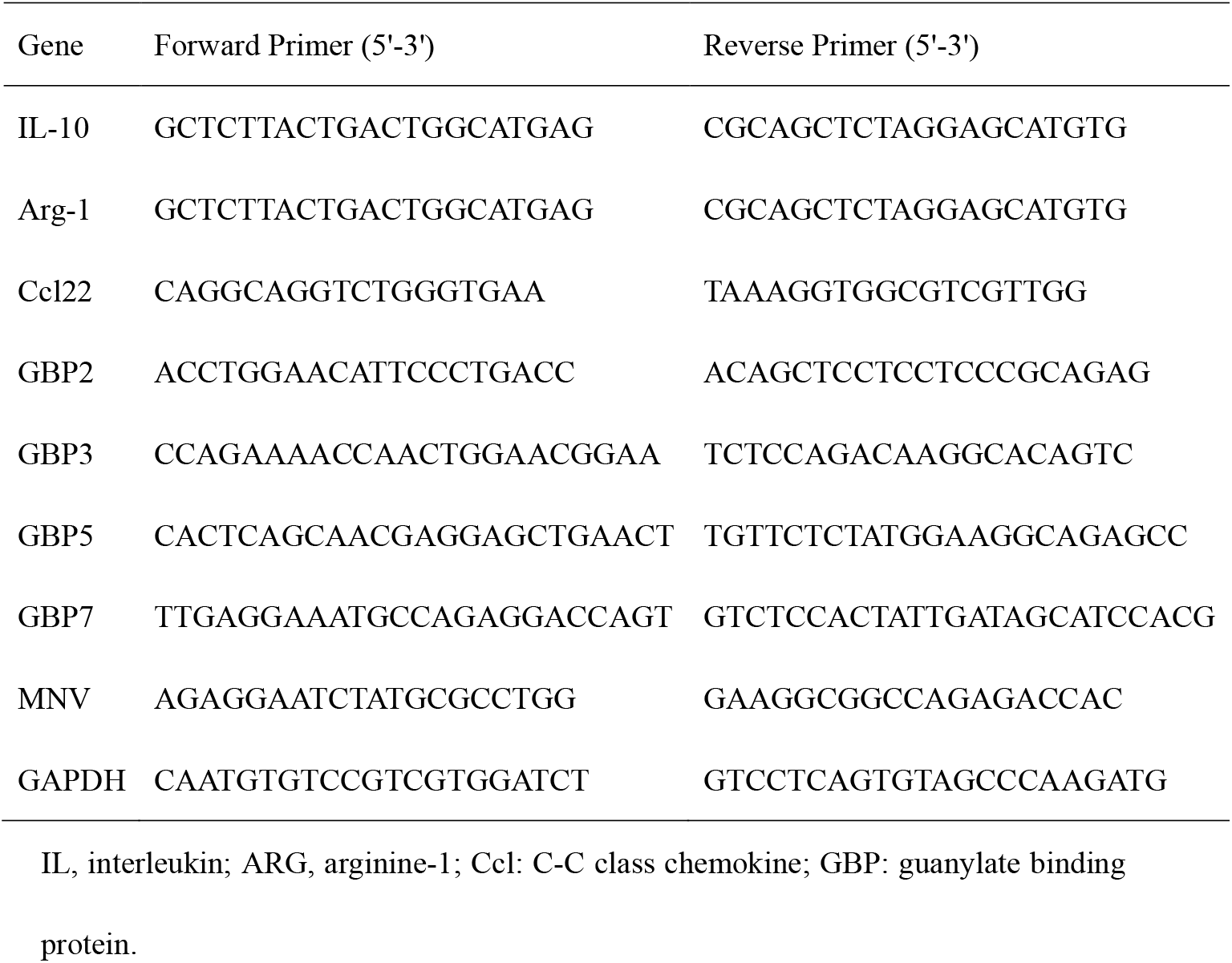
Gene and primer sequences.

### Immunofluore scence assay

An indirect immunofluorescence assay (IFA) for MNV was performed as previously described (Ma, et al., 2020). Briefly, cells were washed with PBS, and fixed with cold acetone/methanol (1:1, v/v) for 20 min at −20°C. The fixed cells were incubated with rabbit anti-MNV VP1 polyclonal antiserum at 37°C for 1 h, washed three times with PBST, and incubated with Alexa Fluor 555-conjugated goat anti-rabbit (Beyotime Biotechnology, Beijing, China) at 37°C for 1 h. After three washes with PBST, infected cells were quantified using a Zeiss LSM510 laser confocal microscope.

### Western blot analysis

Cells were lysed and proteins were separated by 10–15% SDS-PAGE. After 90 min at 120 V, the proteins were electrotransferred to a nitrocellulose membrane (Hybond-C extra, Amersham Biosciences) for 40 min at 20 V. The membrane was blocked with 10% nonfat milk buffer containing 0.1% Tween-20 for 1 h at room temperature, followed by incubation with a rabbit polyclonal anti-VP1 antibody (Ma, et al., 2020) or anti-guanylate binding protein (GBP) 1–5 (Santa Cruz Biotechnology, Santa Cruz, CA, USA). After washing, the membranes were incubated with secondary antibody. The protein bands were visualized using enhanced chemiluminescence detection reagents (Applygen Technologies, Beijing, China). Band intensities were measured by densitometry using ImageJ software (National Institutes of Health, Bethesda, MD). Western blot analyses were repeated three times; β-actin was used for standardization of sample loading. Protein spot levels were determined using ImageJ quantification software.

### RNA library preparation and sequencing

To assess the impact co-incubation of phage vB_SauM_JS25 and MNV on host cell gene expression, RAW 264.7 cells were inoculated with MNV (MOI, 1) at 37°C, and 1 h later, the cells were washed with PBS to aspirate the inoculums and purified phage vB_SauM_JS25 was added to the cells (10^8^ PFU/mL). At 12 h post-phage treatment, the cells were collected. Total RNA was extracted using TRIzol (Invitrogen, CA, USA), and genomic DNA was removed by DNase I digestion (Turbo DNA-free kit; Ambion catalog no. AM1907). RNA integrity was assessed using the RNA Nano 6000 assay kit of the Bioanalyzer 2100 system (Agilent Technologies, CA, USA). Libraries were prepared for sequencing by Cambridge Genomics Services using 3 μg of total RNA and an Ultra^™^ RNA library prep kit for Illumina (NEB, USA). Libraries were quantified by PCR, pooled, and sequenced on an Illumina platform, and 125 /150-bp paired-end reads were generated.

### Data analysis

Raw reads were inspected with FastQC. Reference genome and gene model annotation files were directly downloaded from the genome website. The index of the reference genome was built using Hisat2 (v2.0.5) and paired-end clean reads were aligned to the reference genome using Hisat2. FeatureCounts v1.5.0-p3 was used to count the numbers of reads mapped to each gene. Differential expression analysis of two groups was performed using the DESeq2 R package (1.16.1). DESeq2 provides statistical tests for determining differential expression in digital gene expression data using a model based on the negative binomial distribution. The resulting *P* values were adjusted using the Benjamini–Hochberg approach for controlling the false discovery rate. Genes with an adjusted *P* value < 0.05 found by DESeq2 were considered as differentially expressed. Gene ontology (GO) enrichment analysis of differentially expressed genes (DEGs) was implemented by the cluster profiler R package, in which gene length bias was corrected. GO terms with corrected *P* value < 0.05 were considered as DEGs. To identify and visualize the differential expression of antiviral-related genes between the control (C) groups, phage (P), MNV (M), and MNV plus phage (MP) group, gene expression levels were compared among MP vs M, MP vs P, P vs C, and M vs C using the web tool Venn Diagrams (Hur, et al., 2019).

### Phage DNA transfection assay

Phage DNA was isolated from purified phages (procedure described above) and extracted using a phage DNA isolation kit (ABigen, Beijing, China). Lipofectamine 3000 transfection reagent (Invitrogen, Carlsbad, CA, USA) was used for transient transfection following the manufacturer’s instructions. RAW264.7 cells cultured in 24-well plates were infected with MNV, and then cotransfected with 800 ng of phage gDNA and 100 ng of EGFP plasmid, which served as an internal control. Cell lysates were collected and gene expression levels were detected by RT-qPCR at 24 h post-transfection. Cell viability was tested by CCK-8 assay.

### Confocal microscopy

RAW 264.7 cells were seeded onto 12-mm glass-based dish (Thermo Scientific, USA), allowed to propagate overnight, and subsequently infected with MNV at an MOI of 1. After 1 h, the cells were washed and treated with phages as described above. The cells were fixed with 4% paraformaldehyde for 30 min at 12 h post-phage treatment and permeabilized by incubating in PBS buffer containing 0.2% Triton X-100 for 10 min. The buffer was removed, and the cells were washed three times with PBS. Then staining with primary and secondary antibodies was conducted for 1 h each at room temperature. Primary antibodies against the following proteins were used: MNV VP1 (Ma, et al., 2020) and GBP 1–5. The following secondary antibodies were used: Alexa Fluor 555-conjugated goat anti-rabbit and Alexa Fluor 488-conjugated goat anti-mouse (Beyotime Biotechnology). Samples were washed and labeled with 4’,6’-diamidino-2-phenylindole (Solarbio, Beijing, China) in PBS for 5 min. Glass-based dishe were mounted onto microscope sliders, and the samples were examined using an UltraView Vox confocal microscope (PerkinElmer) (Zhang, et al., 2017).

### Statistical analysis

Data were analyzed for significance by one-way or two-way ANOVA using GraphPad PRISM software (version 5.02 for Windows; GraphPad software Inc.). A *P* value < 0.05 was considered to indicate a statistically significant difference.

## Results

### Inhibitory effect of by bacteriophage vB_SauM_JS25 on MNV multiplication is dose-and time-dependent

To detect the dose–effect relationship between phage vB_SauM_JS25 and MNV replication, RAW 264.7 cells were first inoculated with MNV at an MOI of 1, followed 1 h later by phage vB_SauM_JS25 stimulation at different amounts (10^3^–10^8^ PFU/mL). At 24 h post-stimulation, the cells were lysed and analyzed by RT-qPCR. To investigate the time-dependent suppression of viral replication, cells were seeded in 24-well plates and inoculated with MNV (MOI, 1), followed by vB_SauM_JS25 stimulation at 6, 12, 18, and 24 h post-infection. At 24 h post-stimulation, the cells were lysed and analyzed by RT-qPCR. As shown in **Fig. 1A**, the longer the cells were inoculated with phage, the stronger was the inhibitory effect we observed. The results showed that stimulation with phage vB_SauM_JS25 inhibited MNV replication in a dose-dependent manner (**Fig. 1B**).

**Figure 1.**
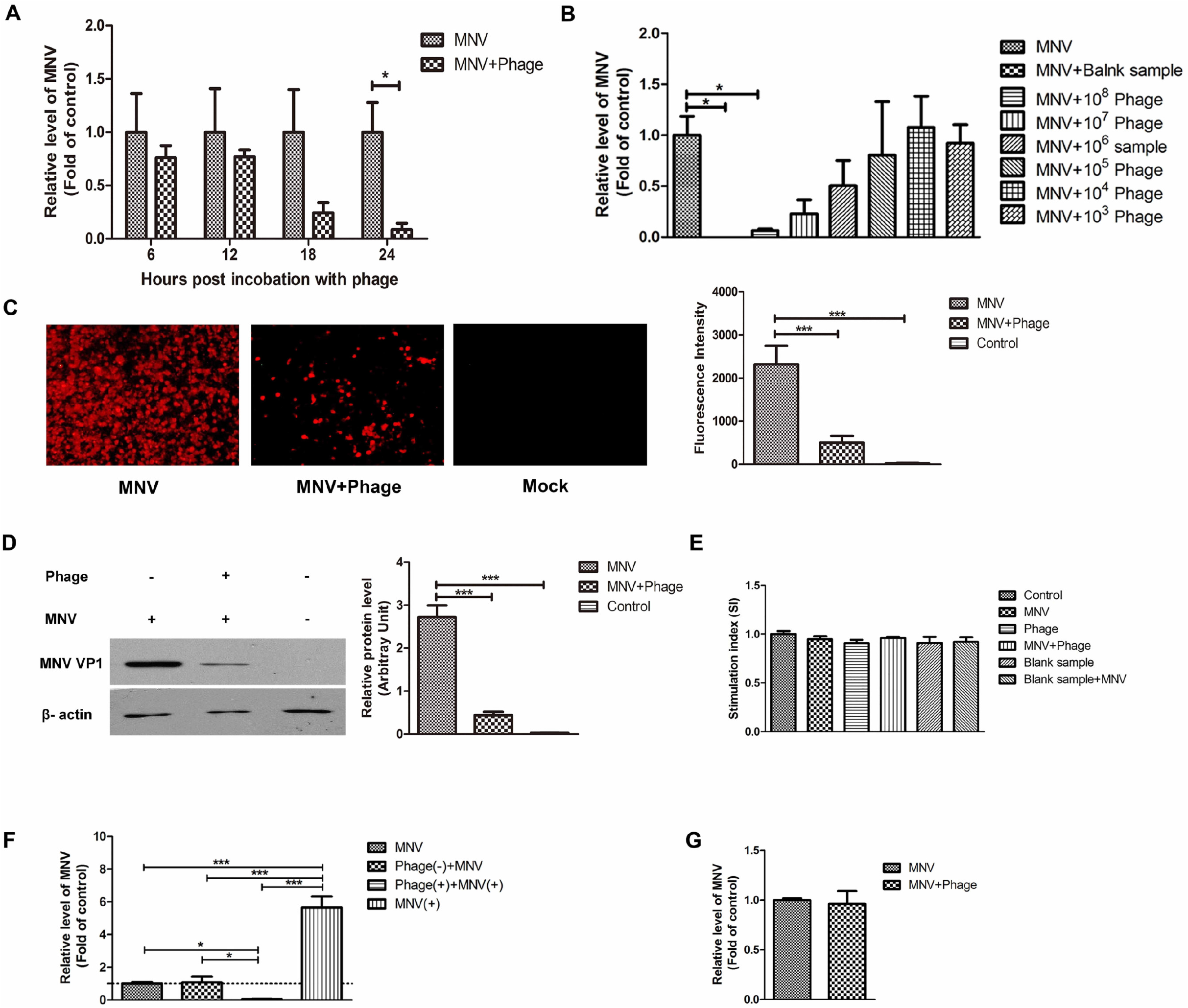
Treatment with phage vB_SauM_JS25 effectively inhibited MNV replication *in vitro*. RAW 264.7 cells were stimulated with MNV (MOI, 1), and 1 h later, cells were challenged with phage. At the indicated time points (6–24 h post-stimulation) **(A)** and at increasing amounts (10^3^–10^8^ PFU/mL) **(B)**, cells were harvested for RT-qPCR. MNV-infected RAW 264.7 cells were stimulated with vB_SauM_JS25 (10^8^ PFU/mL), and 24 h post-stimulation, cells were harvested for indirect IFA **(C)** and western blot analysis **(D)** with anti-MNV VP1. The level of MNV was quantified by immunoblot scanning and normalized to the amount of β-actin. **(E)** Cell viability was assessed by CCK-8 assay. **(F)** RAW 264.7 cells were pre-treated with phage for 3 h and then infected with MNV in the absence (−) or presence (+) of phage for another 24 h. Cells were harvested for RT-qPCR. **(G)** RAW 264.7 cells were infected with MNV for 1 h at room temperature and treated with phage vB_SauM_JS25 (10^8^ PFU/mL) for another 2 h at 37°C. Viral binding of MNV multiplication was analyzed by RT-qPCR. Data are expressed as the mean ± SD (^*^*P* < 0.05, ^**^*P* < 0.01, ^***^*P* < 0.001 compared to the negative control).

Meanwhile, the number of infected cells was determined by western blot and IFA. Fluorescence measurements showed that the number of infected cells was significantly lower (*P* < 0.01) in the MP group compared to the M group (**Fig. 1C**). Similar results were obtained by western blot analysis (**Fig. 1D**). The cytotoxic damage did not differ among the six groups (**Fig. 1E**).

Next, we investigated whether pre-treatment with phage could inhibit MNV replication. RAW264.7 cells were pre-treated with phage for 3 h, and then infected with MNV in the absence (−) or presence (+) of phage for another 24 h. As shown in **Fig. 1F**, consistent treatment with phage significantly decreased the level of MNV mRNA expression (*P* < 0.05). However, when the phage was removed before MNV infection, phage pre-treatment did not influence the MNV replication (*P* > 0.05, **Fig. 1F**). For the viral binding test on MNV of phage vB_SauM_JS25, no significant reduction was observed (*P* > 0.05, **Fig. 1G**).

### RNA-sequencing analysis revealed antiviral effect s of phage vB_SauM_JS25 related with IFN −γ production and response signaling pathways

To further characterize the antiviral effect of phage vB_SauM_JS25 in RAW 264.7 cells, we performed RNA-sequencing (RNA-Seq) to measure the phage–MNV co-incubation samples. RNA-Seq was repeated three times for each group. The following thresholds were used to identify the DEGs between groups: *P* < 0.05 and |log_2_(FC) | ≥ 1 (**Table S1**). A total of 3124 genes were expressed in all four groups (C, P, M, and MP). Venn diagram comparisons for all the expressed genes disclosed the overlap of shared expressed genes among MP vs M, MP vs P, P vs C, and M vs C groups (**Fig. 2**). Between M vs C, 54 DEGs were identified, and between P vs C, 2396 DEGs were identified. Between MP vs P, only 74 DEGs were identified, and between MP vs M, 2399 DEGs were identified, suggesting phage induced most of the response. There were 645 specific genes between MP and M groups, while 1727 genes overlapped between MP vs M and P vs C, indicating that this difference is related with anti-viral effects (Fig. 2). A complete list of DEGs in their respective comparisons is shown in **Table S1**.

**Figure 2.**
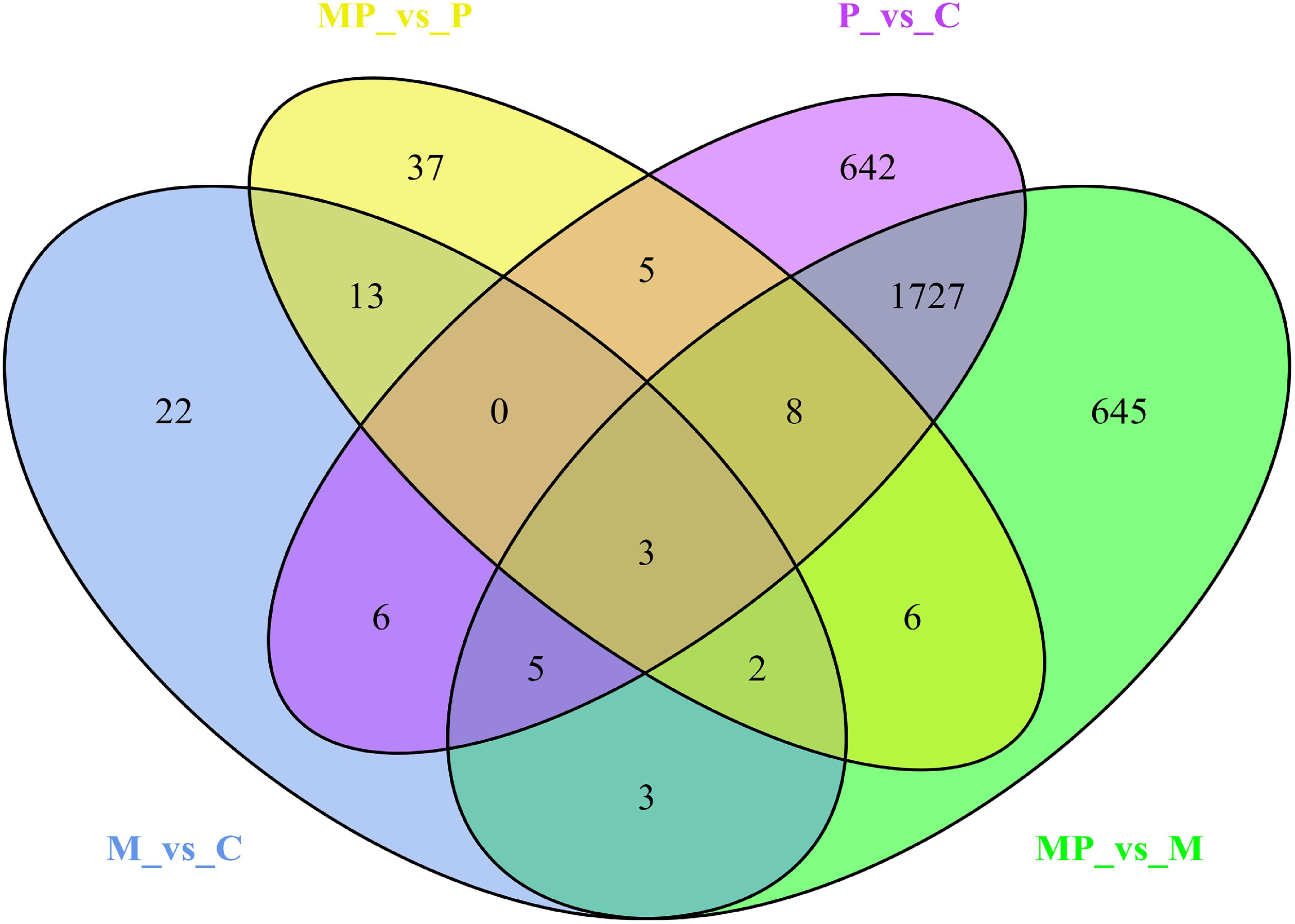
Venn diagram showing the number of differentially expressed genes (DEGs) among different groups. A total of 3,124 genes were expressed in all four groups: control (C), phage (P), MNV (M) and MNV plus phage (MP). The DEGs among MP vs M, MP vs P, P vs C, and M vs C were input to the web tool Venn Diagrams.

GO is a gene function classification system that is often used to describe gene properties. In addition, enrichment analyses of GO terms were performed for DEGs using GOatools. GO terms for which the corrected *P* value < 0.05 were considered significantly enriched (**Table S2**). The top 16 of up-and downregulated DEGs regulating biological processes between the MP and M groups are shown in **Fig. 3**. The upregulated DEGs are predominantly involved in the immune response, immune system processes, and cell adhesion, including the cellular response to IL-1, the cellular response to IFN-γ, and the response to IFN-γ (**Fig. 3**). The downregulated DEGs mainly include genes involved in small molecule metabolic processes and carboxylic acid metabolic processes (**Fig. 3**).

**Figure 3.**
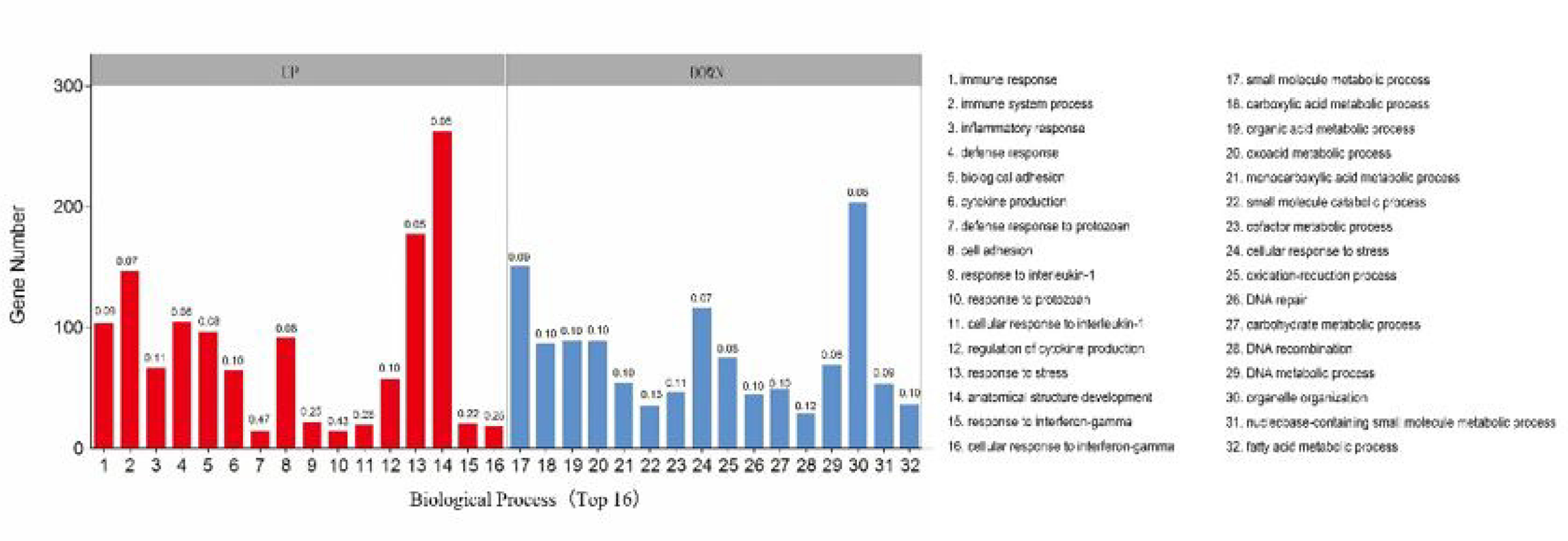
GO enrichment analysis of DEGs between MP and M groups (up- and downregulated genes). *P* values were calculated using the Metacore Tool in the GeneGo package (http://www.genego.com/). GO groups were organized from lowest to highest *P* value. The *x*-axis shows up- and downregulated DEGs with the top 16 enriched GO terms, and the *y*-axis shows the number of DEGs. Enrichment ratios of genes are shown above each column, as well as their GO terms with respect to biological processes.

### Immunity-related GTPases (IRGs) and guanylate-binding proteins (GBPs) were significantly upregulated by phage vB_SauM_JS25

Volcano plots revealed dozens of genes that were upregulated in the MP group compared with M group (**Fig. 4A**). Moreover, phage treatment significantly increased the expression levels of genes related to IFN-γ production and response signaling pathways, which further inhibited MNV replication (**Fig. 4B**).

**Figure 4.**
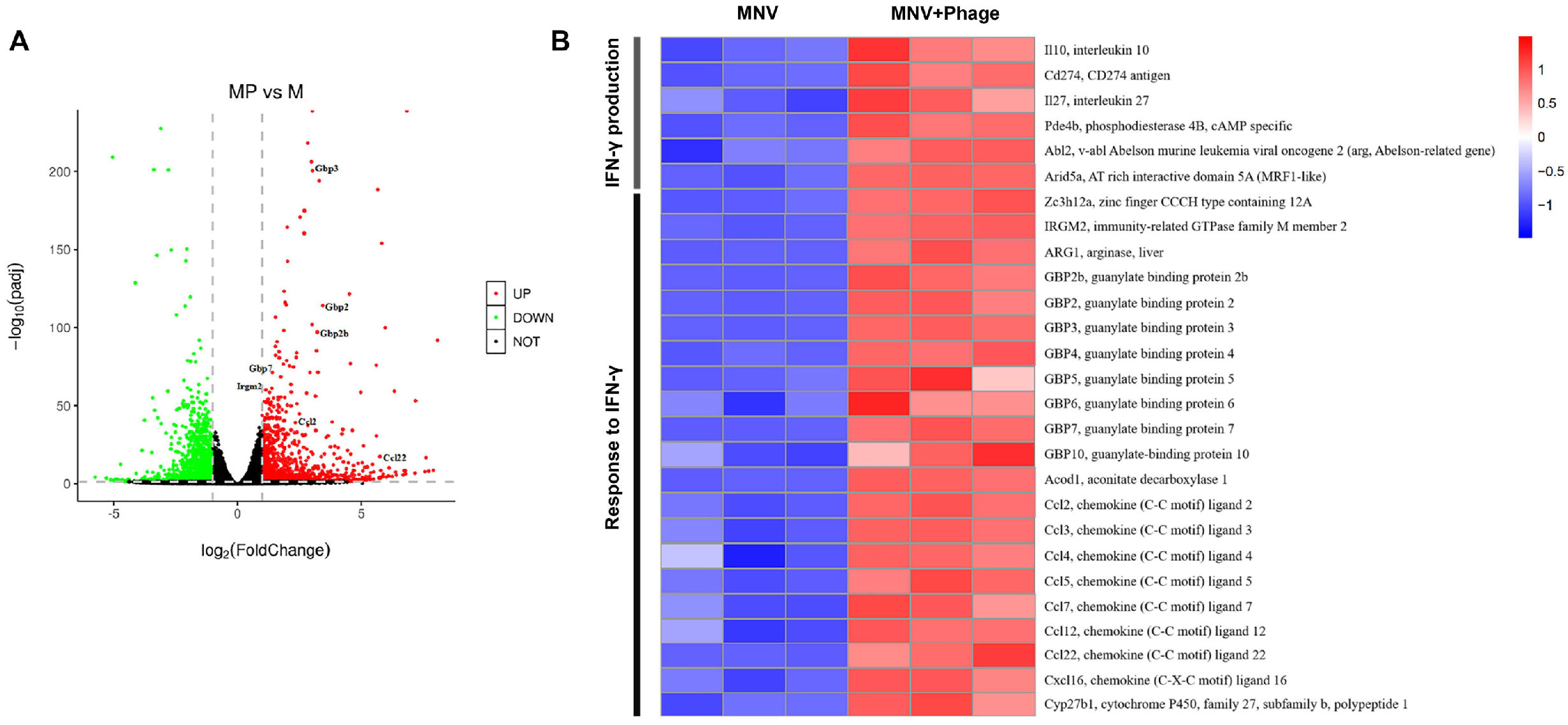
RNA-seq analysis revealed that the antiviral effect of phage vB_SauM_JS25 is related to IFN-γ responses. **(A)** Volcano plot of RNA-seqanalysis in RAW 264.7 cells. **(B)** Heat map of selected gene expression in (A). Gene expression is shown with a pseudocolor scale (from −1 to 1), with the red color indicating a high expression level and the blue color indicating low expression.

In order to validate our RNA-Seq analysis, we selected seven DEGs (*IL-10, Arg-1, Ccl22, GBP2, GBP3, GBP5*, and *GBP7*) and performed RT-qPCR on transcripts from the same biological samples. We observed a strong correlation between and RT-qPCR and RNA-Seq results, confirming the accuracy of the expression data obtained by RNA-Seq (**Fig. 5A–D**). Phage treatment activated the expression of GBPs, as revealed by RT-qPCR analysis, western blot, and confocal microscopy (**Figs. 5D–E and 6A**). Taken together, these data suggested that the IFN-inducible GTPases play an antiviral role against MNV replication *in vitro*.

**Figure 5.**
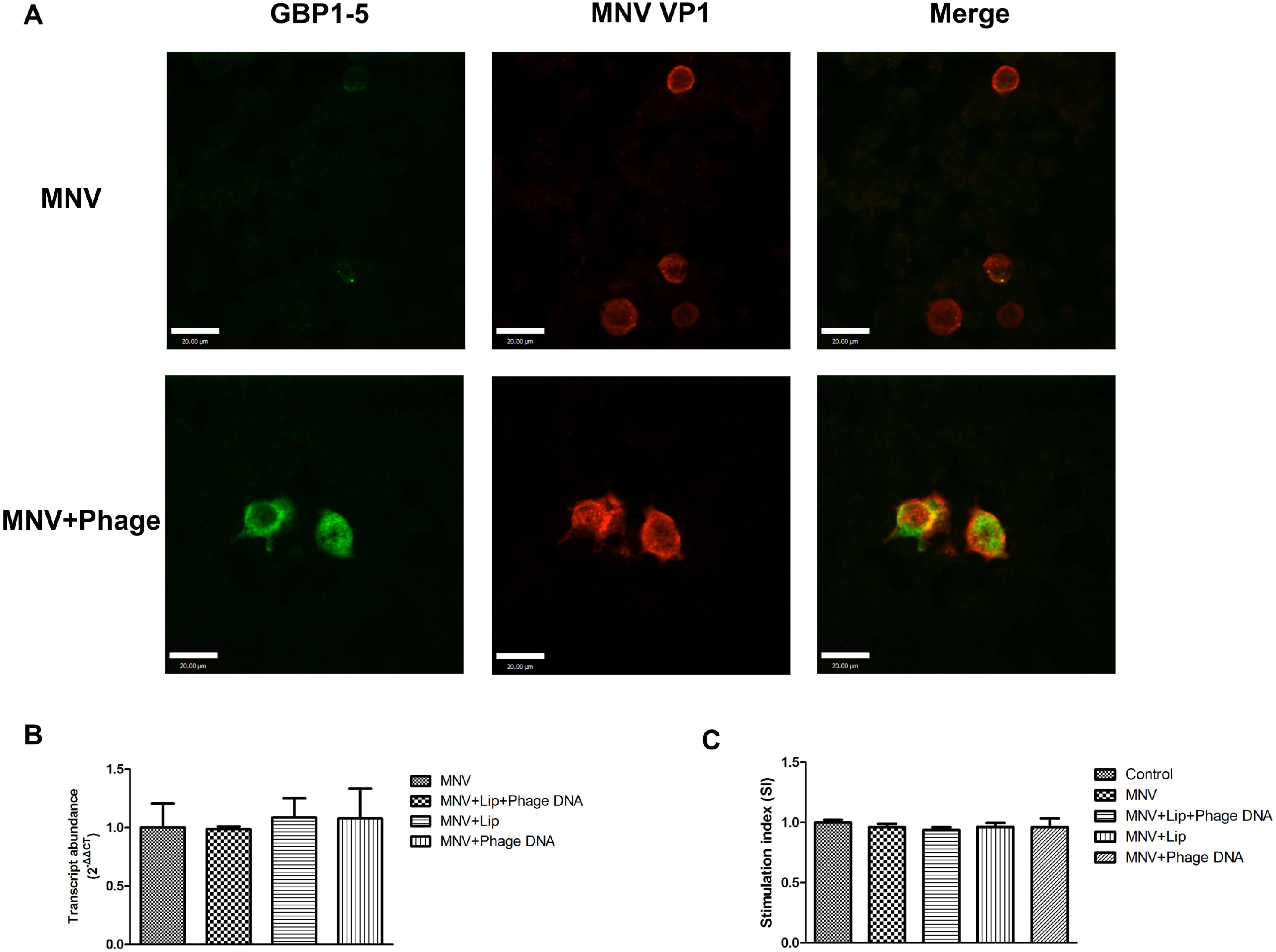
Confirmation of seven upregulated genes *IL-10* **(A)**, *Arg1* **(B)**, *Ccl22* **(C)**, and *GBP2, GBP3, GBP5*, and *GBP7* **(D)** in MNV–phage co-incubation in RAW 264.7 cells by RT-qPCR. *GAPDH* was used as an internal control to normalize the quantitative data. **(E)** MNV-infected RAW 264.7 cells were stimulated with vB_SauM_JS25 (10^8^ PFU/mL), and 24 h post-stimulation, cells were harvested for western blot analysis. The levels of guanylate binding protein (GBP) 1–5 were quantified by immunoblot scanning and normalized to β-actin. Data are shown as mean ± SD (^*^*P* < 0.05, ^**^*P* < 0.01, ^***^*P* < 0.001 compared to the negative control).

**Figure 6.**
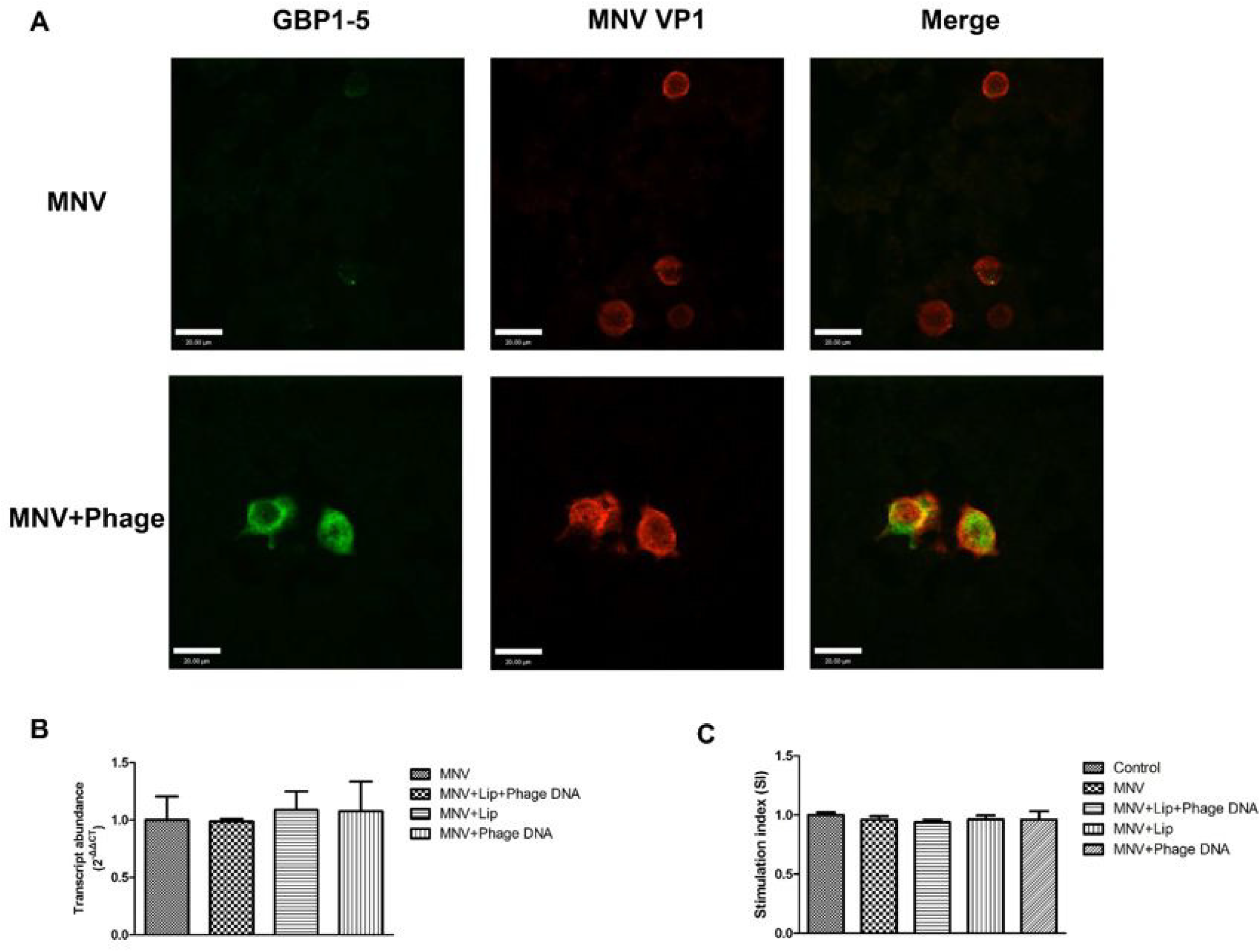
**(A)** GBP1–5 colocalized with MNV VP1 protein in RAW 264.7 cells. Cells were examined by confocal microscopy using UltraView Vox. MNV-infected cells were stained with rabbit anti-VP1 (red) and mouse anti-GBP1–5 (green). Yellow regions are areas of VP1 protein and GBP1–5 colocalization. **(B)** RAW264.7 cells cultured in 24-well plates were infected with MNV and then transfected or stimulated with phage DNA. Cell lysates were collected and analyzed by RT-qPCR at 24 h post-transfection. **(C)** Cell viability was tested in different groups by CCK-8 assay.

To analyze the effects of phage DNA on MNV replication, MNV-infected RAW264.7 cells were transfected or stimulated with phage DNA instead of phage. Phage DNA did not decrease MNV mRNA expression (*P* > 0.05, **Fig. 6B**). Moreover, cell viability was not significantly difference among the five groups (**Fig. 6C**).

## Discussion

The host innate response plays an essential role in the suppression of pathogen infection. The cooperation of phages with the innate immune system was first shown by Tiwari et al. (Tiwari, et al., 2011), who showed the necessity of a neutrophil–phage cooperation in the resolution of *Pseudomonas aeruginosa* infections. This was later confirmed by Roach et al. (Roach, et al., 2017) and Pincus et al. (Pincus, et al., 2015) and an *in silico* model was developed by Leung and Weitz (Leung and Weitz, 2017). It has become clear that phages impact immunity directly, typically through anti-inflammatory effects (Van Belleghem, et al., 2018, Zhang, et al., 2018). Moreover, these phage-induced immune responses could have much broader effects, such as antiviral effects (Sweere, et al., 2019). In this work, we demonstrated the inhibitory effects of phage vB_SauM_JS25 on MNV replication *in vitro*.

In this study, we demonstrated *Staphylococcal* enterotoxins could be efficiently removed from phage lysate and free of endotoxin contamination. Hence, the immune responses observed are induced by the phages. In the present study, transcriptomes of RAW 264.7 cells inoculated with MNV and phages were sequenced using an Illumina platform. A total of 3124 genes were expressed in all four groups (C, P, M, and MP). Comparison of P vs C groups identified 2396 DEGs, while only 74 DEGs were identified between the MP vs P groups, suggesting that the phage itself induced most of the response. Between the MP vs M groups, 2399 DEGs were identified. Moreover, of the DEGs between MP vs M groups, 645 specific and 1727 genes overlapped between MP vs M and P vs C, indicating that these DEGs are related with antiviral effects. In addition, enrichment analyses of GO terms were performed for DEGs using GOatools between MP and M groups (Klopfenstein, et al., 2018). We observed a strong correlation between RT-qPCR and RNA-Seq results, confirming the accuracy of the expression data obtained by RNA-Seq. The upregulated DEGs were predominantly involved in the immune response, immune system processes, and cell adhesion, especially the cellular response to IFN-γ and the response to IFN-γ, to which MNV replication is sensitive. The downregulated DEGs were mainly involved small molecular metabolic processes, carboxylic acid metabolic processes, and small molecule catabolic processes. Although Gogokhia et al (2019) reported that phages could activate IFN-γ by stimulation of TLR9 (Gogokhia, et al., 2019), no similar activation was observed in our transcriptomics data. The phage DNA showed no antiviral activity after transfection or stimulation in RAW264.7 cells. Thus, the antiviral effects of phages might not be mediated by TLR9, but might rather be based on other undiscovered mechanisms.

Immunity-related GTPases (IRGs) and guanylate binding proteins (GBPs), upon their induction by IFN-γ, are known to oligomerize in a GTP-dependent manner on the membrane of pathogen-containing vacuoles, leading to vesiculation and subsequent disruption of vacuolar structures (Pilla-Moffett, et al., 2016). Similarly, IRGs and GBPs may disrupt the structural integrity of the MNV replication complexes (Biering, et al., 2017). Moreover, we found that IRGs and GBPs, including *IRGM2, GBP2b, GBP2, GBP3, GBP4, GBP5, GBP6, GBP7*, and *GBP10*, were significantly upregulated by phage vB_SauM_JS25 in RAW 264.7 cells. IRGs and GBPs are dynamin-like GTPases that are targeted to the membrane of cytoplasmic vacuoles containing bacteria or viruses (Ferreira-da-Silva Mda, et al., 2014, Biering, et al., 2017). Given this, the data shown here indicate that the expression of both IRGs and GBPs was induced by phages, reflecting their direct antimicrobial effects on the membrane structure of the MNV replication complexes. GBPs and IRGs induced by phages may contribute to clearance of bacteria and viruses, revealing a synergy with host innate immunity. However, the roles of IRGs and GBPs in mechanisms underlying phage-induced antiviral effects should be further studied.

Human norovirus is a major cause of nonbacterial gastroenteritis worldwide. Due to the lack of robust cell culture and small animal systems, little is known about human norovirus pathogenicity. Murine norovirus (MNV) is used as a model to study norovirus infection. Disruption of the gut microbiome by human norovirus reported in a previous human study suggested that the gut microbiome is closely related to the immune response to norovirus (Nelson, et al., 2012). It has been demonstrated that MNV infection stalls host protein translation and the production of antiviral and proinflammatory cytokines (Fritzlar, et al., 2019). This evasion strategy delays innate immune responses to MNV infection and accelerates disease onset. In this study, the immune response to IFN-γ, including the expression of specific cytokines and chemokines, such as IRGM2, also referred to as chemokine (C-C motif) ligand 2 (CCL)2), was involved in the immune response against norovirus infection. Therefore, the data indicate that phages could activate the antiviral response and exert a positive effect on human cells.

Overall, our results show that phage vB_SauM_JS25 could significantly inhibit MNV replication in a time- and dose-dependent manner, but these inhibitory effects did not occur during the binding stage of MNV to RAW 264.7 cells. RNA-Seq was performed to investigate the interaction between phages and cells. We found that the upregulated DEGs were mainly involved in the immune response, immune system processes, and cell adhesion, especially the cellular response to IFN-γ and the response to IFN-γ. Volcano plots revealed IRGs and GBPs were upregulated in the MP group. To our knowledge, this study is the first to show the inhibitory effects of phages on MNV by activating GBPs. Further studies involving genetically engineered phages and *in vivo* experiments should be performed to evaluate which components of phages are responsible for their antiviral activity. This work provides new insight into the interplay between phages and eukaryotic cells, establishes a general understanding of phage-based antiviral effects, and paves the way for the development of antiinfection agents.

## Acknowledgments

This work was funded by the National Natural Science Foundation of Jiangsu Province (BK20180054), the National Natural Science Foundation of China (Grant number 31602078), and the Jiangsu Agricultural Science and Technology Foundation (CX(19)2014).

